# Learning to See Again: Biological Constraints on Cortical Plasticity and the Implications for Sight Restoration Technologies

**DOI:** 10.1101/115188

**Authors:** Michael Beyeler, Ariel Rokem, Geoffrey M. Boynton, Ione Fine

## Abstract

The “bionic eye” – so long a dream of the future – is finally becoming a reality with retinal prostheses available to patients in both the US and Europe. However, clinical experience with these implants has made it apparent that the vision provided by these devices differs substantially from normal sight. Consequently, the ability to learn to make use of this abnormal retinal input plays a critical role in whether or not some functional vision is successfully regained. The goal of the present review is to summarize the vast basic science literature on developmental and adult cortical plasticity with an emphasis on how this literature might relate to the field of prosthetic vision. We begin with describing the distortion and information loss likely to be experienced by visual prosthesis users. We then define cortical plasticity and perceptual learning, and describe what is known, and what is unknown, about visual plasticity across the hierarchy of brain regions involved in visual processing, and across different stages of life. We close by discussing what is known about brain plasticity in sight restoration patients and discuss biological mechanisms that might eventually be harnessed to improve visual learning in these patients.

## 2. INTRODUCTION

The field of sight restoration has made dramatic progress in the last five years. The Argus II device (epiretinal, Second Sight Medical Products Inc., Rizzo et al. 2014, da Cruz et al. 2016) as well as the Alpha-IMS system (subretinal, Retina Implant AG, Stingl et al. 2015) recently completed clinical trials and are now available for commercial sale in the US and Europe. Several other electronic vision implants have either started or are planning to start clinical trials in the near future. These include the IRIS II (epiretinal, Pixium Vision, Hornig et al. 2017), PRIMA (subretinal, Stanford University, Lorach et al. 2015), as well as devices by the Bionic Vision Australia consortium (suprachoroidal, Ayton et al. 2014, Saunders et al. 2014), Nidek Co. Ltd. (suprachoroidal, Fujikado et al. 2016) and Nano Retina (epiretinal). At the same time, sight recovery technologies based on optogenetics and gene therapies are also making considerable strides; over a dozen human gene therapy trials are underway (Petrs-Silva and Linden 2014), and optogenetic clinical trials will likely begin within the next two years (Busskamp et al. 2012). Within a decade, many individuals suffering from blindness are likely to be offered a wide range of options for sight restoration that depend on widely different technologies (Fine et al. 2015, Ghezzi 2015).

Clinical trials suggest that, for some patients, visual prostheses can provide visual information that is useful in daily life; enabling these individuals to perform simple object localization, motion discrimination, and letter identification tasks (Zrenner et al. 2011, Humayun et al. 2012, Stingl et al. 2013, Dagnelie et al. 2017). However, these reports also highlight the limitations of current devices. Clinical and psychophysical measurements of patients’ experiences have made it apparent that the vision provided by current devices differs substantially from normal sight. Only a handful of patients show performance close to the theoretical limits that would be expected if performance were limited by the spacing and density of the electrodes. Insights from psychophysical experiments and theoretical considerations (reviewed in Fine and Boynton 2015) suggest that this may be due to interactions between implant electronics and the underlying neurophysiology of the retina, discussed in more detail in Section 3 below. In electronic and optogenetic devices these include unselective stimulation of retinal cells (Figure 1B), and in the case of epiretinal devices, spatial blurring due to axonal comets (Panel C). In the case of optogenetic technologies this includes temporal blurring (Panel D).

**Figure 1.**
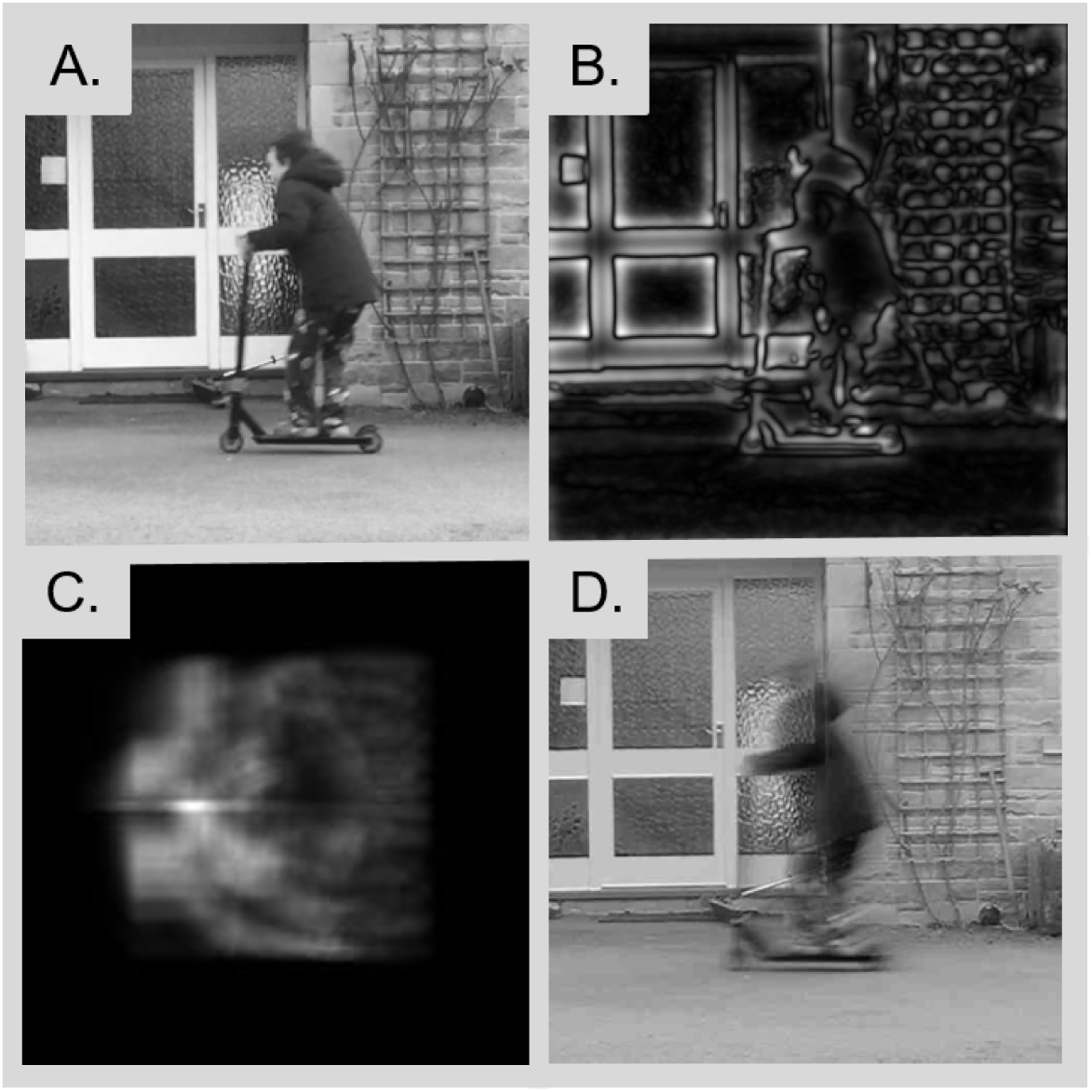
Example models demonstrating potential perceptual distortions and information loss for different sight recovery technologies. All images are based on the central 12° region of a movie and a 1000x1000 array (A.) Original image. (B.) Simulation of sub-retinal electrical stimulation, based on simultaneously stimulating ON and OFF pathways with no axonal stimulation. (C.) Simulation of epiretinal electrical stimulation, based on simultaneously stimulating ON and OFF pathways with axonal stimulation resulting in visual ‘comets’. (D.) One frame in a movie showing simulation of optogenetic stimulation, based on a model of simulating ON pathways with an optogenetic protein with temporal dynamics based on ReaChR (Sengupta et al. 2016). Motion streaks are the result of the sluggish temporal dynamics of ReaChR. Modified figure based on Fine and Boynton (2015).

Patients report the experience of prosthetic vision as being like *“looking at the night sky where you have millions of twinkly lights that almost look like chaos”* (Pioneer Press 2015). One particular challenge is dealing with the combination of a limited field of view and an external camera, which can be pointed in a direction other than the patient’s direction of gaze (Barry and Dagnelie 2016). As a result, patients are required to take in the world in a piece-wise fashion, using mental imagery to connect individually perceived phosphenes to form a whole. One patient reported: *“I do see boundaries. I can create an image. I know what a car should look like. I know what a tree should look like. I know what houses should look like. I know what objects should look like, and I have those images as memories in my brain. With this device, you can start to create an image by going back and forth, checking the boundaries and borders. We don’t have natural saccade movement helping us. The camera is right above my nose in a fixed position. It doesn’t move. You can create an image, but it takes a lot of time and a lot of work.”* (Discovery Eye Foundation 2014).

Consequently, it might be more appropriate to think of current retinal prostheses as ‘providing visual input’ rather than ‘restoring sight’. Nonetheless, it is the hope of the sight restoration community that the artificial visual signal elicited by electrical prostheses can help patients in daily life. One critical open question is – what role will cortical plasticity play in helping patients make use of this artificial visual input? Previous experience with cochlear implants suggests that cortical plasticity is capable of compensating for significant loss of information in the sensory input. Although the auditory signal from the earliest cochlear implants was too diminished to allow for speech perception (Eisen 2003), current implants can mediate surprisingly good speech recognition in most adult-deafened subjects (Shannon 2012). Patients successfully adapt to the impoverished and distorted auditory input over the course of hours to approximately a year.

Plasticity for retinal implants may be very different than for cochlear implants. Early stages of the visual hierarchy seem to be much less plastic than the auditory or somatosensory systems: the preponderance of evidence, reviewed below, suggests that large-scale cortical reorganization at the level of the lateral geniculate nucleus (LGN) or the primary visual cortex (V1) may be very limited. This lack of plasticity early in the visual processing stream has important implications for what expectations are reasonable when hoping that patients can learn to interpret the input from visual prosthetics.

The goal of the present review is to briefly summarize the vast basic science literature on perceptual learning and cortical plasticity with an emphasis on how this literature might relate to the field of visual prostheses. The objective is not to provide concrete recommendations for prosthetic development or rehabilitative training, but to provide insight into the fundamental principles which govern perceptual learning and cortical plasticity throughout the lifespan, so as to inform the choices of engineers and clinicians who are designing devices or training patients.

We begin with a description of the distortions and information loss likely to be experienced by visual prosthesis users. We then discuss cortical plasticity and perceptual learning and describe what is known, and what is unknown, about visual plasticity; both across the hierarchy of brain regions involved in visual processing, and across different stages of life. Because we assume that most visual prostheses will be implanted in late blind individuals suffering from binocular loss (at least in the foreseeable future), certain important fields in the cortical plasticity literature are neglected in this review. These include the effects of stroke or trauma within subcortical or cortical structures, monocular deprivation (for reviews see Levi et al. 2015, Kiorpes 2016), and the development of cross-modal plasticity (for review see, Bavelier and Neville 2002, Lewis and Fine 2011) since this tends to be far more limited in late than early blind individuals (for discussion see Collignon et al. 2013). We close by discussing what is known about brain plasticity in sight restoration patients and discuss biological mechanisms that might eventually be harnessed to improve perceptual learning.

## 3. DISTORTIONS AND INFORMATION LOSS IN PROSTHETIC DEVICES

As mentioned in the Introduction, interactions between implant electronics and the underlying neurophysiology of the retina will differ depending on the device. Nonselective stimulation of retinal cells is a concern for all electronic and optogenetic technologies. Electrode sizes of current prosthetic devices (typically on the order of 100 μm) inevitably lead to the indiscriminate stimulation of thousands of morphologically distinct retinal cells (including simultaneous activation of both ON and OFF cells, Figure 1B). This is in contrast to natural stimulation, which precisely activates a number of specialized, functionally complementary, parallel processing pathways in the retina (for a review see Nassi and Callaway 2009).

While using large electrodes exacerbates the issue, nonselective stimulation remains a concern for electronic technologies with extremely small electrodes and optogenetic technologies. In optogenetics, while it is possible to target a single class of cell (e.g. bipolar vs. ganglion cells) it is not currently possible to selectively target ON vs. OFF cells (see Fine et al. 2015, for a review). Thus, for both these technologies, the lack of differentiation in neuronal responses across different cell types results in a reduction in the amount of information available to cortex (Moreno-Bote et al. 2014). Prostheses implanted in the LGN and cortex will similarly suffer from an information-limiting lack of differentiation across the stimulated neural population, with the exact effects depending on which layers/regions of cortex are implanted.

There are of course many other causes of perceptual distortions and information loss. For epiretinal prostheses a significant concern are the visual “comets” (Figure 1C) that result from axonal stimulation. For optogenetic technologies, there is likely to be loss of temporal resolution, due to the sluggish temporal dynamics of optogenetic proteins (e.g. ReaChR). As is observable in Figure 1D, this loss of *temporal* resolution can result in significant spatial distortions for rapidly moving objects. Finally, the distortions and information losses for subcortical and cortical prostheses are likely to be complex, and depend significantly on which layers (Pezaris and Reid 2007) and/or cortical areas are stimulated. Direct electrical stimulation of the LGN (Pezaris and Reid 2007) or early visual cortex (V1/V2) elicits phosphenes (Tehovnik and Slocum 2007, Murphey et al. 2009). In contrast, electrical or magnetic stimulation of areas like the middle temporal visual area (V5/MT, Beckers and Homberg 1992), inferotemporal cortex, or specific regions within the fusiform gyrus (Afraz et al. 2006, Rangarajan et al. 2014) can disrupt motion and face processing, respectively, but does not elicit percepts (Murphey et al. 2009, Rangarajan et al. 2014). Representations in these higher-level areas may occur at a distributed network level (Levy et al. 2004). As a consequence focal stimulation does not create an interpretable percept, but can disrupt the network activity elicited by a natural visual stimulus.

Because each type of device is likely to suffer from unique types of information loss, quantitative measures of the quality of visual information provided by a device may become critically important as patients are given choices across multiple very different technologies. Similarly, being able to assess the ‘visual cost’ of a given distortion could play an important role in prosthetic design. There are already numerous metrics that can assess the perceptual quality of an image or movie based on models of the visual system (Haines and Chuang 1992, Wang et al. 2004, Laparra et al. 2016), but these assume no capacity for plasticity. Metrics describing the amount of ‘neurophysiologically available’ information (i.e. assuming perfect plasticity) in a device are therefore also important. Figure 2 describes one way in which loss of information (in this example resulting from axonal stimulation in an epiretinal prosthesis) might be quantified. A variety of devices with the number of electrodes ranging from two to 1500 were simulated. The field of view was held roughly constant at 10x20 degrees of visual angle and the electrode radius was set to 20% of the electrode-electrode spacing. Spatial distortions due to axonal stimulation was predicted using a model based on Nanduri et al. (2011). The resulting ‘predicted percepts’ (Panels A and B) for each device were used as input into a principal components analysis (PCA). If every electrode acted independently, they would be able to span a space of possible percepts with a dimensionality equal to the number of electrodes in the array. We can quantify the actual dimensionality of the space by the number of principal components needed to explain 95% of the variance – this can be thought of as the number of independently acting, ‘effective’ electrodes. Panels C and D show the number of effective electrodes as a function of the number of physical electrodes in the device both without (Panel C) and with (Panel D) axonal stimulation. Without axonal blurring, the number of principal components increases linearly with the number of electrodes with a slope of roughly 0.5: in an array of 1400 electrodes about 700 principal components are required to explain 95% of the variance. When axonal stimulation is included the number of effective electrodes drops by about an order of magnitude; with 1400 electrodes, around 50 principal components are required to explain 95% of the variance.

**Figure 2:**
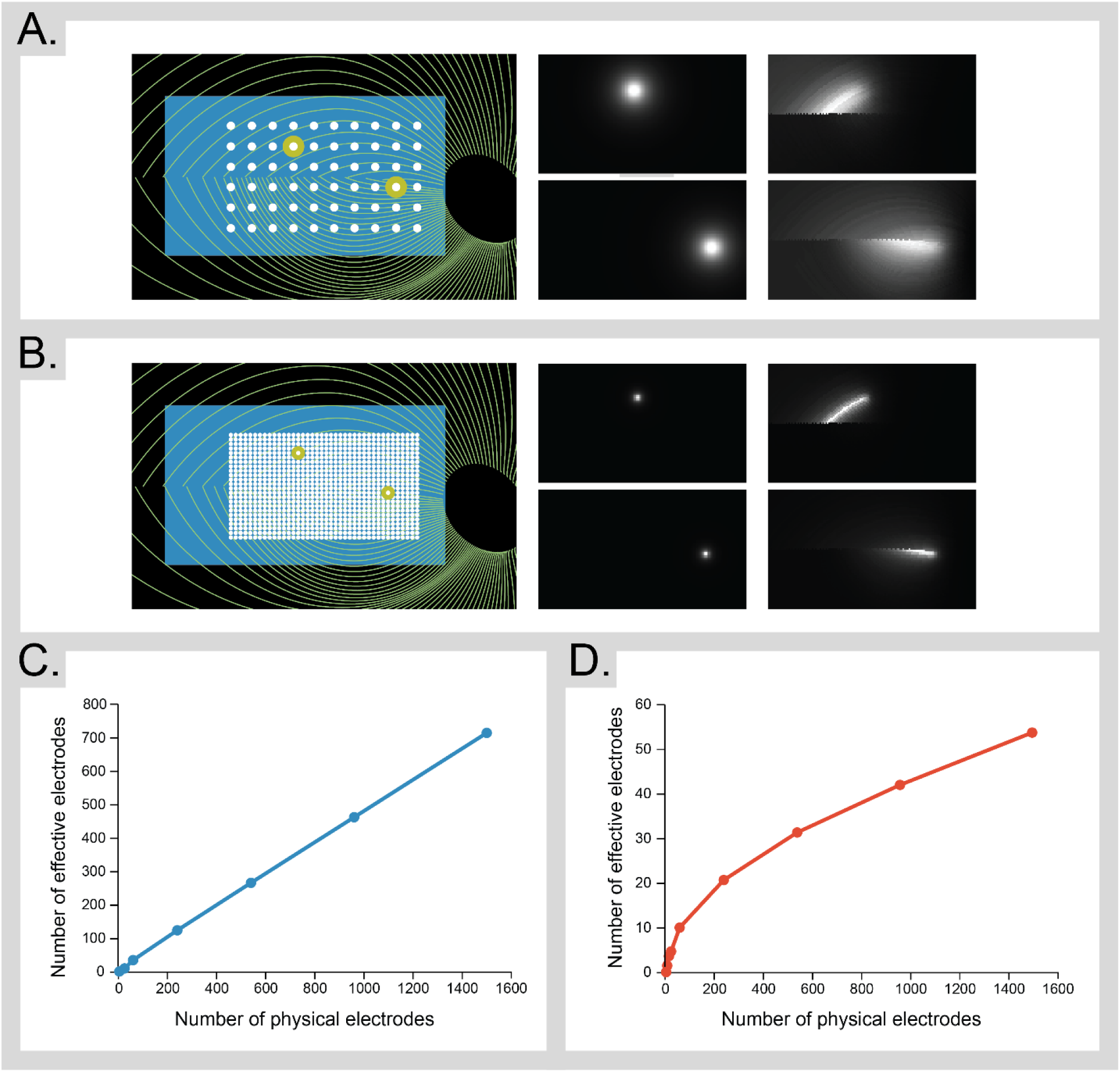
Quantifying loss of information in epiretinal devices. (A.) The retinal location of a simulated Argus II array (6x10 electrodes) and the predicted percepts generated by stimulating a single electrode at a time both without (left) and with (right) axonal stimulation. (B) The retinal location of a simulated high-resolution array (30x50 electrodes) with analogous percepts. (C, D.) The number of effective electrodes (i.e., the number of principal components needed to explain 95% of the variance) as a function of the number of physical electrodes in the array, with (C.) and without (D.) axonal stimulation.

## 4. CORTICAL PLASTICITY AND PERCEPTUAL LEARNING: TWO SIDES OF THE SAME COIN

Cortical plasticity (also known as neuroplasticity) is an umbrella term that describes the ability of cortex to change its structure or function in response to experience. Cortical plasticity can be observed at multiple temporal scales (Horton et al. 2017), ranging from short-term (seconds to minutes) to long-term (days to many months). Cortical plasticity also occurs across a wide range of spatial scales, ranging from alterations in the tuning characteristics of individual neurons up to reorganization of entire neuronal circuits (“cortical remapping”, see Wandell and Smirnakis 2009, for a review).

The behavioral manifestation of cortical plasticity is *perceptual learning*. In a traditional perceptual learning paradigm, training on a specific task leads to a long-lasting improvement in behavioral performance (Karmarkar and Dan 2006, Deveau et al. 2014), such as a decrease in the minimal orientation difference that can be detected, or an increase in the speed for detecting target shapes embedded in distracters (for a review, see Sagi 2011). Similar to cortical plasticity, perceptual learning is a well-described feature of mammalian visual systems that occurs over multiple time scales – ranging from seconds (Mooney 1957) and minutes to years (Krupinski et al. 2013).

Although perceptual learning and cortical plasticity represent corresponding behavioral and physiological measurements of the same phenomenon, it has often proved difficult to understand the relationship between the two. One reason for this is that performance improvements for apparently quite similar tasks can be mediated by very different cortical substrates. For example, training in auditory (Recanzone et al. 1993) or tactile (Recanzone et al. 1992a, Recanzone et al. 1992b) frequency discrimination seems to result in “cortical recruitment” within primary sensory areas; that is, expansion of the amount of cortical territory/number of neurons representing the trained frequencies. In contrast, training on visual orientation discrimination tasks does not seem to substantially alter either the number of neurons tuned to the trained orientation or the orientation tuning of neurons whose orientation preference is close to the trained frequency (Crist et al. 2001, Ghose et al. 2002; though see Schoups et al. 2001 for evidence of subtle changes in tuning). Rather, visual perceptual learning is linked to alterations in extra-classical contextual responses (Crist et al. 2001, Li et al. 2004), possibly mediated by altered response properties within V4 neurons with orientation tuning relevant to the task (Yang and Maunsell 2004).

## 5. PERCEPTUAL LEARNING

Perceptual learning is likely to be very different in prosthesis users than in sighted subjects. In subjects with normal vision it can be assumed that the organization of the visual pathways is already close to optimal, especially in the fovea (Westheimer 2001). As a result, task performance for naturalistic tasks such as object recognition is generally at ceiling, making it necessary to titrate difficulty by reducing stimulus difference (e.g. orientation difference), contrast or stimulus duration, or by adding external noise. However, even if the task is made more difficult, learning is likely still limited by the fact that the visual system is already optimized for these stimuli - indeed this is probably why familiar objects show so little learning (see task 6, Figure 3, below).

**Figure 3:**
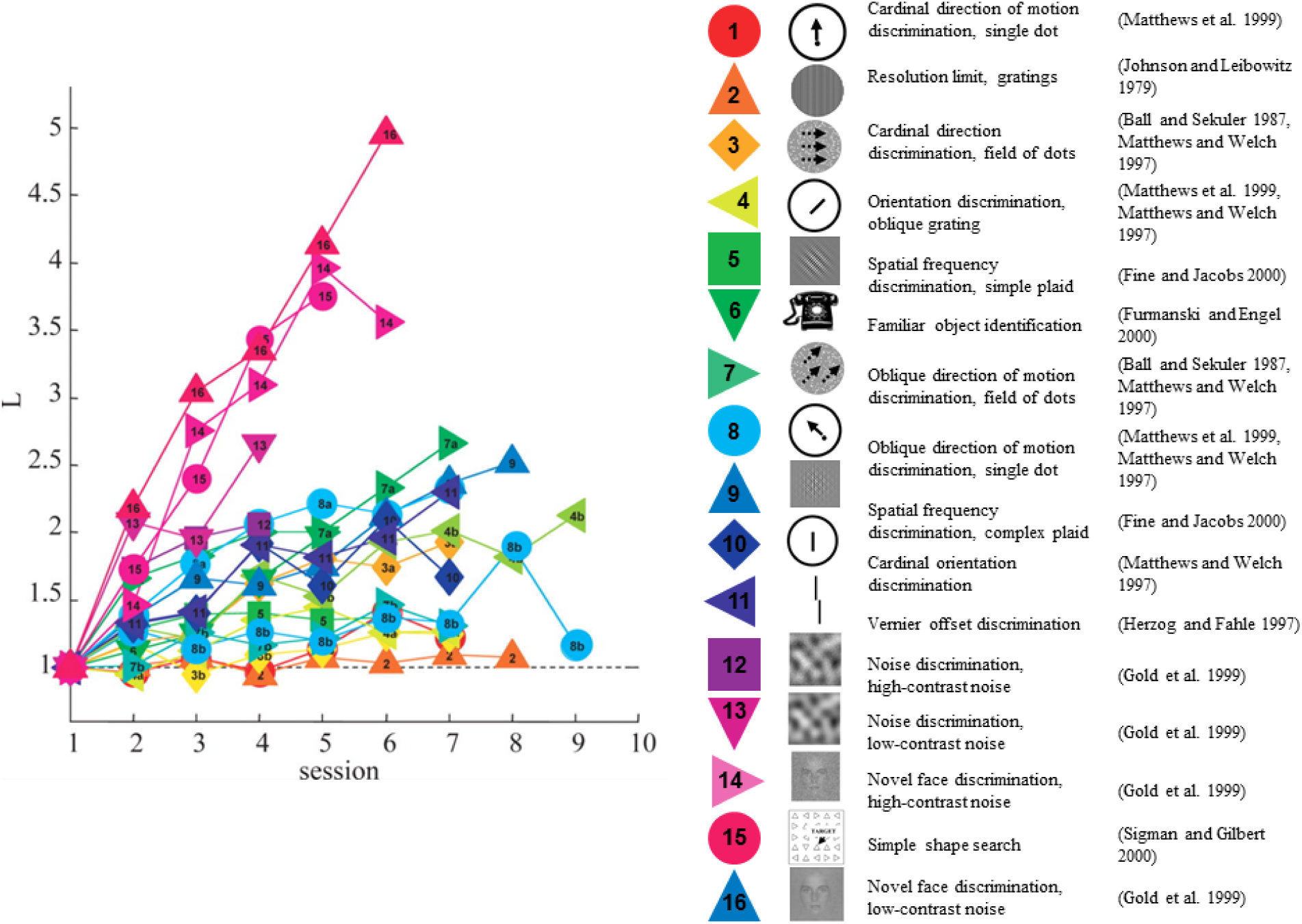
Performance improvement as a function of practice over time for 16 different perceptual tasks (ordered according to the estimated learning slope). For all studies, performance on each session was converted into d’ (Green and Swets 1966). The learning index, L, measures improvements in performance with practice, 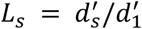, where s is the session number. A learning index remaining near 1 implies that observers showed no improvement in performance with practice. Reprinted with permission (Fine and Jacobs 2002).

### 5.1. Learning as a function of stimulus complexity

As shown in Figure 3, the speed and extent of learning varies dramatically across different kinds of tasks or stimuli. While it is possible to produce significant improvements in performance for tasks performed on basic attributes of a stimulus that are represented in V1, such effects tend to require very extensive training over up to tens of thousands of trials over many days or weeks. Examples include grating detection (e.g. Li et al. 2008, Yehezkel et al. 2016), orientation tuning (Fine and Jacobs 2000), contrast discrimination (Dorais and Sagi 1997, Adini et al. 2002, Yu et al. 2004), and visual acuity (Landolt C and two-line resolution thresholds). In contrast, learning for more complex higher-order properties occur more rapidly, with a few sessions and several hundred trials of practice (Fine and Jacobs 2000), including discriminations that are dependent on context (such as three-line bisection, Vernier discrimination, texture discrimination, contour detection, and shape discrimination).

Similar findings exist in the clinical literature. For example, reading speed can improve significantly with practice in patients with advanced macular degeneration (Chung 2011, Astle et al. 2015, Maniglia et al. 2016). However these improvements seem to be mainly mediated by a reduced susceptibility to crowding rather than an improvement in low-level acuity (Chung 2013, He et al. 2013).

One possible explanation for larger learning effects in more complex tasks is that low-level representations may be resistant to plasticity in adulthood. Another possibility is that the larger amount of perceptual learning observed for complex tasks or stimuli may reflect the cumulative effects of changes in tuning properties across the visual processing cascade. Both possibilities are discussed in more detail below.

### 5.2. Generalization and specificity in perceptual learning

The goal of training with prosthetic stimulation is to improve visual performance outside the laboratory. Thus, it is critical that learning is generalizable: improvements in performance must transfer to untrained stimuli or tasks. In general, the transfer of learning from one task to another only occurs when the two tasks share cognitive elements (Woodworth and Thorndike 1901). However, other aspects of training also play a critical role. Improvement for difficult tasks (Ahissar and Hochstein 1997, Talluri et al. 2015), and/or tasks involving simple features, such as contrast detection (Swift and Smith 1983) or discrimination (Dorais and Sagi 1997, Yu et al. 2004) tends to be highly specific. Improvements fail to generalize to untrained tasks performed with the same stimuli, or within the same task to untrained spatial frequencies, orientations, retinal locations, contrasts, or even eye of origin.

It is important to note that specificity in learning for properties like orientation or spatial location does not necessarily imply that learning occurs within early cortical regions, where contrast, orientation, and spatial invariance has not yet been achieved. Rather, learning may be mediated by the retuning of higher-level neurons to become more specific for these properties (Mollon and Danilova 1996, Petrov et al. 2005, Zhang et al. 2010).

Consistent with the perceptual learning literature on sighted subjects, patients implanted with retinal prostheses show barely any improvement on simple perceptual tasks, such as contrast sensitivity (Castaldi et al. 2016) or motion discrimination (Dorn et al. 2013, Castaldi et al. 2016), even after extensive training over several months. The limited and highly specific perceptual plasticity observed for low level features such as contrast or motion direction has important implications for visual prosthetic design. For example, electronic prostheses have been shown to be fairly limited in the number of luminance levels (typically between four and 16, depending on the task and implant) that can be discriminated (Greenwald et al. 2009, Stingl et al. 2015). Similarly, current optogenetic technologies for restoring retinal function are expected to have a limited dynamic range for luminance (Gaub et al. 2015, Sengupta et al. 2016). In both technologies, sensitivity to contrast is also likely to be compromised by the absence of light/dark adaptation mechanisms within photoreceptors and horizontal cells (Bownds and Arshavsky 1995). The psychophysical literature described above suggests that it could be overly optimistic to expect patients to show improved sensitivity to low-level properties such as contrast as a result of experience or training, except under highly specific contexts unlikely to be of much use in real-world vision.

With more complex tasks, making stimuli variable and the task easier (Ahissar and Hochstein 1997, Talluri et al. 2015) has been shown to aid generalization. In the absence of stimulus variability, learning remains highly specific – generally at the most specific level possible (Ahissar and Hochstein 2004). But under conditions where stimuli vary widely generalized learning can occur, e.g., when presented at multiple spatial locations or with multiple different exemplars, or when stimulus differences are relatively large, and performance levels are high (Xiao et al. 2008, Wang et al. 2012, Wang et al. 2014, Baeck et al. 2016).

One important observation for visual prosthetic technologies is that real-world tasks that require higher level representations, such as object recognition, might not require compensation for the distortions of prosthetic vision to occur at the earliest stages of processing (such as in V1). For example, while it may be impossible to learn to see more brightness levels, it may be possible to develop higher level representations of objects that are less susceptible to this loss of grayscale information. For example, training with two-tone (Mooney) faces has been shown to improve face/non-face discrimination abilities (Latinus and Taylor 2005).

Finally, the nature of feedback should be considered carefully. Whenever possible, feedback should not be binary (correct/incorrect), since this gives patients very little information about the nature of their error. Rather, it would be more effective to give patients auditory or tactile information about how large their error was, and in what direction. This approach has been shown to improve performance in a target localization task (e.g. Barry and Dagnelie 2016, Endo et al. 2016).

In summary, many of the discoveries from the perceptual learning literature on sighted individuals provide ‘guidance principles’ that will be helpful in developing effective training paradigms for prosthetic or optogenetic patients. For example, although an individual using a prosthetic device would likely show far more perceptual learning than sighted subjects for a grating orientation task, this would be predicted from the basic science literature: this task for a prosthetic patient is probably much more like a complex shape discrimination task, and learning increases with stimulus complexity. The other principles of effective learning, as described above, are also likely to influence rehabilitation protocols for prosthetic devices. For example, training on a very specific task (e.g., discriminating two particular orientations) may not transfer well to very different orientations or spatial frequencies. Similarly, the finding from the basic science literature that working at too difficult a performance level impedes learning (DeLoss et al. 2014) is also likely to apply to individuals struggling to interpret prosthetic input. Current training procedures for prosthetic users have tended to focus on maintaining a high performance level. As a result, training has relied on simple tasks with restricted stimulus sets or a limited number of navigational environments. Such task designs might have a fairly limited potential for generalizing beyond laboratory settings. Better rehabilitation designs are likely to involve patients learning to make relatively coarse discriminations using a broad stimulus set in the context of an easy task.

### 5.3. Training and ‘gamification’ within virtual environments

Designing rehabilitation paradigms for prosthetic patients that consist of easy tasks involving coarse discriminations across broad stimulus sets has proved difficult, since using a large stimulus set also tends to make the task much more difficult for patients. In contrast to sighted people whose perceptual performance is already excellent, a patient with prosthetic vision must learn to adjust to any or all of the following: camera/eye-position disconnection (Barry and Dagnelie 2016), limited field of view, poor contrast, spatial distortions that are not constant across the visual field, a luminance-brightness relationship that varies across the retina, and percepts that rapidly fade over time.

It is already recognized that “isolating skills and using materials that are designed to support the development of these skills” (Dorn et al. 2013) is an important aspect of training for prosthetic device users. The instructional kit provided by Second Sight Medical Products Inc. consists of high-contrast items of simple shape (e.g., plates and bowls) presented on a black background. According to Ghodasra et al. (2016), some investigators have added items of intermediate contrast to the training set. However, for the reasons described above, training with such a restricted stimulus set may not lead to generalizable learning. Moreover, for tasks outside object identification, such as navigation, it is extremely laborious and expensive to create environments that allow for task simplification (e.g. dark rooms with white doors).

This difficulty in creating simple navigation environments has led to the suggestion that patients be trained in a familiar environment (Ghodasra et al. 2016). However, evidence from the basic science literature suggesting that learning generally occurs at the most specific level possible raises the concern that learning to navigate a specific environment may not generalize to unfamiliar environments. There is also the concern that learning to use distorted visual information to navigate in a familiar environment for which there is a rich internal memory representation may be a very different process from using that distorted information to make sense of an unfamiliar environment.

‘Virtual environments’, which have recently been successfully used for training with sensory substitution devices (Maidenbaum et al. 2013, Maidenbaum et al. 2014, Chebat et al. 2015, Levy-Tzedek et al. 2016, Maidenbaum et al. 2016), offer an elegant way to solve many of these difficulties. Within the VR context it is easy to generate a varied stimulus set of objects or environments. Direct (i.e., not via the camera) stimulation of electrodes can be used to create a tailored environment that can gradually and systematically shape adaptation to the various abnormalities of prosthetic vision across a wide range of tasks. Tasks and environments can be specifically designed to master accommodation to one aspect of prosthetic vision (e.g. using head motion to explore the visual field) before moving to the next (e.g. contrast), and so on. It is possible to gradually increase task complexity (e.g. by gradually increasing the number of objects to be recognized, gradually moving towards realistic contrast levels, or gradually increasing the complexity of the environment to be navigated) while keeping performance at a high accuracy level so as to maximize generalization. Virtual environments can also allow for continuous haptic or auditory feedback – for example using the frequency of a tone to tell subjects that they are drifting off course in a navigation task, or haptic feedback to shape hand-eye co-ordination by gradually ‘narrowing’ the target window within which a subject can successfully grasp an object.

A final advantage of virtual environments is that they allow for ‘gamification’ - embedding the task within a video game context. It has recently been discovered that learning, especially for simpler features that are normally resistant to generalizable learning, can be enhanced through gamification (e.g. Li et al. 2011, Deveau et al. 2014). Gamification has been shown to improve perception, visuomotor coordination, spatial cognition, and attention; with effects remaining even two years after the end of intervention (Feng et al. 2007, Bavelier et al. 2010).

There are a number of reasons why the gaming context may be particularly effective for inducing plasticity: perceptual training within a gaming context generally includes more varied stimuli/surround context than standard experimental perceptual learning regimes, and might recruit top-down, attentional systems, possibly altering the excitatory/inhibitory balance to allow for heightened plasticity (Feng et al. 2007), as described in more detail below. Gamification may also be particularly effective at recruiting reward and attentional neuromodulators such as dopamine (Koepp et al. 1998) and acetylcholine (Bavelier et al. 2010, Rokem et al. 2010, Rokem and Silver 2013) since video games are undoubtedly more engaging than standard psychophysical lab experiments. Finally, asking subjects to perform a task that is intrinsically rewarding is more likely to be successful in engaging patients in self-guided training – thereby reducing the need for personalized rehabilitation therapy.

## 6. CORTICAL PLASTICITY WITHIN THE SENSITIVE PERIOD

Although implantation of prosthetic implants will be restricted to adults for the foreseeable future, the literature on plasticity in the sensitive period is nonetheless highly relevant. Important aspects of visual cortical architecture, such as retinotopic organization, are driven by molecular signaling and are robust to visual perturbations even during early development. It is important to recognize that aspects of the visual architecture that are minimally dependent on experience during early development are similarly unlikely to be modifiable via training in adulthood.

### 6.1. V1 cortical plasticity

Early in postnatal development many tuning properties within primary visual cortex are remarkably plastic. Visual experience plays an important role in shaping the formation of ocular dominance columns, the size and orientation of receptive fields, as well as spatial frequency, direction, and disparity tuning of V1 neurons (Wiesel and Hubel 1963, Wiesel and Hubel 1965, Raviola and Wiesel 1978, Movshon and Van Sluyters 1981, Sherman and Spear 1982, Ackman and Crair 2014). In the extreme, in animal models, when visual inputs are redirected into the auditory thalamus, the auditory cortex remodels to process visual information (reviewed in Horng and Sur 2006).

In contrast to most other tuning properties, visual experience plays only a minor role in the development of cortical retinotopic organization, which is primarily driven by molecular signaling (Huberman et al. 2008, Cang and Feldheim 2013). In the mouse, cortical retinotopic organization persists in the absence of retinal waves, albeit with reduced precision (Grubb et al. 2003, McLaughlin et al. 2003, Cang et al. 2005). Similarly, in the macaque, adult-like connections between V1 and V2 are present before LGN axons reach layer IV (Coogan and Van Essen 1996), although further refinement occurs with the onset of visual experience (Barone et al. 1995, Batardiere et al. 2002, Baldwin et al. 2012).

In humans, disruption of visual experience in infancy seems to have limited effects on cortical retinotopic organization. The retinotopic organization of corticocortical connections between V1-V2-V3 can be observed on the basis of BOLD resting state correlations in both early blind and anophthalmic (in which both eyes fail to develop) individuals (Bock et al. 2015, Striem-Amit et al. 2015). Rod monochromats virtually lack cone photoreceptor function and therefore have a retinal scotoma within the all-cone foveola. In these individuals some of the V1 region that normally responds to the foveola responds to rod-initiated signals originating from neighboring regions of the retina (Baseler et al. 2002), suggesting some reorganization, but this organization is limited in spatial extent, and may be mediated by an expansion of receptive fields rather than by a fundamental reorganization of the retinotopic map.

The predominance of molecular cues in determining retinal organization is also observable in individuals born without an optic chiasm or even with only one cortical hemisphere. Individuals with albinisms (Hoffmann et al. 2003, Hoffmann et al. 2012), FHONDA syndrome (Ahmadi et al. 2016) or born with one cortical hemisphere (Muckli et al. 2009) have a miswiring of the retinal-fugal projection. Because the retinotopic organization of cortex is primarily determined by molecular cues rather than visual experience this miswiring results in overlapped cortical representations of left and right visual hemifields where each region of cortex represents two distant (mirror symmetric) locations in visual space (Hoffmann et al. 2003, Hoffmann et al. 2012).

### 6.2. Plasticity beyond V1

In humans, there is a variety of evidence for remarkable flexibility in decoding congenitally abnormal V1 representations. For example, as described above, in individuals suffering from a miswiring of the retinal-fugal projection the retinotopic organization within V1 is highly abnormal. Despite this, acuity losses in these individuals are dominated by their foveal hypoplasia, and they show no perceptual confusion across mirror symmetric locations in the two visual hemifields (Bao et al. 2015). This suggests that a strikingly abnormal retinotopic organization within V1 can be successfully perceptually decoded by later stages of visual processing.

PD, an individual who had central cataracts resulting in annular pupils until the age of 43 similarly showed a host of perceptual adaptations to his distorted visual input, including suppression of diplopic images, and enhanced gain control for spatial frequencies that were heavily attenuated by his poor optics (Fine et al. 2002a). Similarly, individuals with complete Schubert–Bornschein CSNB1 genetic deficits are thought to have severely compromised on-bipolar pathways (Cibis and Fitzgerald 2001, Bijveld et al. 2013). Yet these patients show surprisingly good visual performance under photopic conditions, with an average visual acuity of 0.3 logMAR (Zeitz et al. 2015) and report no perceptual difficulties beyond their acuity loss (M. Neitz 2015, personal communication).

In animal models, relatively few studies have examined the effects of congenital visual loss on higher levels of visual processing. However, as described above, when visual inputs are redirected into the auditory thalamus it not only elicits remodeling of the auditory cortex (reviewed in Horng and Sur 2006), but these responses can also guide simple visual behaviors, suggesting that visual information from auditory cortex can be functionally integrated into the visual pathways (von Melchner et al. 2000).

## 7. ADULT PLASTICITY

One important question is whether adult plasticity in visual cortex differs from juvenile plasticity quantitatively, or whether there are also qualitative differences (Karmarkar and Dan 2006). Specifically, as described below it is not yet clear whether or not adult visual cortex can generate novel, functionally appropriate synaptic connections.

### 7.1. V1 plasticity

Since the classic study of Wiesel and Hubel (1965) it has been clear that the plasticity of early visual areas declines sharply in adulthood. Kittens who had one eye sutured shut from birth until three months of age did not have functional vision in the closed eye, even after the eye had been reopened for a considerable period. This deterioration of functional vision did not occur in cats deprived in adulthood, even when the eye was sewn shut for an entire year, showing that the effects of deprivation were far less extreme after the end of the critical period.

Curiously, tactile and auditory sensory areas seem to retain considerably more plasticity in adulthood than visual areas. Both tactile (Recanzone et al. 1992a, Recanzone et al. 1992b) and auditory (Recanzone et al. 1993, Ohl and Scheich 2005, Polley et al. 2006) primary sensory areas have been demonstrated to show dramatic reorganization in adult animals. The reason for the relative lack of plasticity in V1 as compared to auditory or tactile sensory areas remains unclear. One possible explanation is that there is significantly more subcortical processing within somatosensory and auditory than within visual pathways. Thus, A1 and S1 can be considered as ‘higher’ in their respective processing pathways than V1 is within the visual processing hierarchy. If so, the lack of plasticity in V1 is consistent with the idea that plasticity increases across the sensory hierarchy (Hochstein and Ahissar 2002, Ahissar and Hochstein 2004), see Section 7.3.

Two main types of experimental paradigms have been used to measure cortical plasticity in V1: training on feature-based tasks that vary in their complexity, and the effect of retinal lesions.

#### 7.1.1 The effect of feature-based training on V1 responses

The most common task that has been used to assess adult V1 plasticity is orientation discrimination. While one monkey electrophysiology study found subtle changes in orientation tuning after several months of practice on an orientation discrimination task (Schoups et al. 2001), this result was not replicated by Ghose et al. (2002) using a very similar paradigm. Electrophysiology studies using more complex tasks such as dot (Crist et al. 2001) or line bisection (Li et al. 2004) similarly did not find alterations in basic receptive field properties in V1, but did find that training altered top-down contextual influences. One possibility is that effects of learning for orientation within V1 are predominantly due to contextual modulation of extra-classical receptive field properties of V1 responses (Gilbert et al. 2001) and/or modifications in feedback responses into V1 (Sagi 2011). This interpretation is consistent with observations that (1) the effects of perceptual training on neuronal responses vanish when the trained task was not performed or when the monkey was anesthetized (Li et al. 2004), (2) experiments probing contextual effects, such as in contour integration, seem to display stronger effects than tasks like simple orientation discrimination (Gilbert et al. 2001), and (3) perceptual learning is tightly linked with attention (Crist et al. 2001).

Using fMRI, studies in human V1 have found evidence of some plasticity in adult visual cortex: enhanced responses to trained orientations have been found after training with a grating detection task (Furmanski et al. 2004) and an orientation texture discrimination task (Schwartz et al. 2002). Training in orientation discrimination also produced more discriminable cortical responses (Jehee et al. 2012). However, because of the sluggish hemodynamic response measured with BOLD imaging, it is not clear whether these alterations are driven by changes in bottom-up receptive field properties or top-down responses. Although one EEG study (Pourtois et al. 2008) found, in contrast to these previous studies, that training *reduced* V1 responses starting as early as 40ms post-stimulus, this was an exploratory study that has yet to be replicated.

#### 7.1.2 The effect of retinal scotomata on V1 responses

Several studies have found that retinal lesions in the cat induce extremely rapid axonal sprouting and pruning in cortical areas corresponding to the scotoma (Darian-Smith and Gilbert 1994, Yamahachi et al. 2009, Marik et al. 2014). However, it is not clear whether this represents ‘reawakening’ of functional juvenile plasticity or more closely resembles the corruptive retinal remodeling that occurs in late stages of photoreceptor disease (Marc and Jones 2003).

Several studies examining cortical functional tuning after retinal lesions in cats and primates have reported that neurons in the lesion projection zone (LPZ) became responsive to light stimulation of the retina surrounding the damaged area (Kaas et al. 1990, Gilbert and Wiesel 1992, Darian-Smith and Gilbert 1995, Abe et al. 2015) consistent with large-scale reorganization within cortex. However, other studies failed to find evidence of cortical reorganization after retinal lesions (Rosa et al. 1995, Horton and Hocking 1998, Smirnakis et al. 2005).

Similarly, some human fMRI studies have shown responses in the LPZ (Baker et al. 2005, Baker et al. 2008, Dilks et al. 2009) or shifts in measured cortical retinotopic organization (Ferreira et al. 2017) in late blind individuals suffering from visual field loss due to retinal dystrophies, while other studies have failed to find evidence for reorganization (Masuda et al. 2008, Masuda et al. 2010, Baseler et al. 2011).

These discrepancies between reported outcomes in both the neurophysiology (also see Calford et al. 2005, Smirnakis et al. 2005, for alternative explanations) and the fMRI literature may possibly be explained by many studies including one of two possible confounds: (1) using a model to estimate cortical retinotopic organization that does not consider the absence of input when the stimulus was in the scotoma, and/or (2) failing to account for top-down attentional effects As far as the modeling confound is concerned, failing to factor in the absence of input when the stimulus falls within the scotoma biases receptive field estimates in a way that closely mimics the expected effects of cortical reorganization (Binda et al. 2013, Haak et al. 2015). All the neurophysiological (Kaas et al. 1990, Gilbert and Wiesel 1992, Darian-Smith and Gilbert 1995, Abe et al. 2015) and fMRI (Baseler et al. 2011, Ferreira et al. 2017) studies that *did* find shifts in retinal organization in V1 used methods susceptible to this modeling confound. Indeed Baseler et al. (2011) found dramatic shifts in cortical retinotopic organization in late blind macular degeneration patients. However, they did *not* attribute these shifts to plasticity because almost identical shifts were observed in normally sighted individuals with simulated scotoma (Baseler et al. 2011, Binda et al. 2013).

As far as the attentional confound is concerned, robust responses to visual stimuli are found in the lesion projection zone when subjects perform a one-back task (Baker et al. 2005, Baker et al. 2008, Dilks et al. 2009) but not during passive viewing (Rosa et al. 1995, Horton and Hocking 1998, Smirnakis et al. 2005, Masuda et al. 2008, Masuda et al. 2010, Baseler et al. 2011), suggesting that the signals observed are driven by a top-down mechanism, rather than reorganization of the bottom-up sensory signal.

In summary, neurophysiological (Rosa et al. 1995, Horton and Hocking 1998, Smirnakis et al. 2005, Masuda et al. 2008, Masuda et al. 2010, Baseler et al. 2011) and fMRI (Rosa et al. 1995, Horton and Hocking 1998, Smirnakis et al. 2005, Masuda et al. 2008, Masuda et al. 2010, Baseler et al. 2011, Shao et al. 2013) studies that were robust to both of these two confounds have uniformly failed to find retinotopic reorganization within cortex.

A second difficulty in the literature is that, because of the lack of a retinal dystrophy model in primate, animal studies have relied on experimenter-induced retinal lesions, rather than the gradual loss of function over time that occurs in patients with retinal dystrophies. One study, that used a primate identified as having a condition that approximated juvenile macular dystrophy, found no evidence of recruitment of the lesion projection zone (Rosa et al. 1995, Horton and Hocking 1998, Smirnakis et al. 2005, Masuda et al. 2008, Masuda et al. 2010, Baseler et al. 2011, Shao et al. 2013).

Thus, compensation for the distortions and information loss of visual prosthetic devices is likely to rely on plasticity in visual cortical areas central to V1.

### 7.2. Plasticity beyond V1

#### 7.2.1 The effects of training and altered visual experience

The neurophysiology literature suggests that in adult animals basic tuning tuning properties in higher-order areas of the visual cortex are more malleable with training than those in V1. For example, in contrast to V1, orientation tuning in V4 does change after training in an orientation discrimination task (Yang and Maunsell 2004). Analogous shaping of neuronal tuning to match task demands has also been found for more complex stimuli within primate inferotemporal cortex, the highest level of the ventral visual stream. When trained with shape stimuli that varied across four features, an enhanced neuronal representation was found for features that were important for the categorization task, relative to features that were irrelevant to the task (Sigala and Logothetis 2002). Similarly, training monkeys to discriminate novel visual stimuli causes the emergence of a population of IT neurons which respond selectively to these novel stimuli (Kobatake et al. 1998, DiCarlo and Maunsell 2000), or which become capable of distinguishing between them (Jagadeesh et al. 2001).

In humans, the effects of altered visual experience on higher-level vision comes from case-studies of patients that have had their sight ‘restored’ after prolonged vision loss by ophthalmological procedures such as cataract (Fine et al. 2002b, Ostrovsky et al. 2009, Sinha et al. 2013, McKyton et al. 2015) or corneal replacement surgery (Fine et al. 2003b, Sikl et al. 2013). Improvements in the contrast sensitivity function has been noted in several (though not all) individuals who had sight restored at a young age (between the ages of 8-17 years) (Kalia et al. 2014). This ability to show learning for the contrast sensitivity function may be age–dependent: improvements were not found in MM (Fine et al. 2003a) or PD (Fine et al. 2002a, a case of recovery from low vision). Both MM and PD had their sight restored in their 40s.

Impairments in shape processing, object recognition, and face processing are also observed in sight recovery patients, and these persist even after more than a decade of restored optical sight (Huber et al. 2015). Interestingly these deficits are observable both in individuals who have sight restored in late childhood (Sinha et al. 2013) and in a patient, KP, who lost vision at 17, and had his sight restored at age 71 (Sikl et al. 2013). Thus, the sensitive period for deprivation appears to be broader (extending to at least well into the teenage years) than the critical period for recovering normal vision. Interestingly, motion processing, including shape from motion, seems to be relatively robust to the effects of prolonged visual deprivation (Fine et al. 2003b, Saenz et al. 2008). In individuals who have suffered from prolonged blindness, visual prosthetics may prove to be more effective in providing input that can mediate ‘dorsal-stream’ tasks such as navigation (which are also less reliant on higher spatial frequencies), than ‘ventral-stream’ tasks such as reading, face recognition or object recognition.

#### 7.2.2 Can higher-level visual areas compensate for losses earlier in the pathway?

Only one study has specifically examined plasticity in accessing patterns of activity within V1. Ni and Maunsell (2010) examined the effect of prolonged training on detection thresholds for microstimulation within V1 of the macaque. Although it was possible to train the animals to become experts at detecting microstimulation of visual cortex, this expertise came at the cost of impaired detection of visual stimuli at the same retinotopic location. Interestingly, this effect was reversible with retraining on detecting the visual stimulus – suggesting that the local circuitry (whether within V1 itself, or within a higher cortical area decoding V1 activity) somehow reconfigured to better detect task-relevant patterns of neuronal activity.

In humans, a common clinical example of long-term adult-onset distortion of V1 representations is macular degeneration. One of the first symptoms of this disease (earlier even than the presence of a noticeable blind spot), is visual distortion. Typically, straight lines appear wavy or crooked, and the aspect ratio of objects is distorted (Gerrits and Timmerman 1969, Kapadia et al. 1994, Safran et al. 1999, Safran et al. 2000, Zur and Ullman 2003, Dilks et al. 2007, Mavrakanas et al. 2009). One likely explanation for these perceptual distortions is that shifts or expansions in receptive field properties within early visual areas, such as V1, are not fully compensated for in later decoding (Dilks et al. 2007).

The classic experimental paradigm for examining the effects of distorted visual input are prism experiments (e.g., Stratton 1897a, Stratton 1897b, Kohler 1951). Subjects asked to wear optical prisms that produce a misalignment between visual and proprioceptive information initially show biases toward the virtual (perceived) target position (for a recent review, see Sachse et al. in press). After more prolonged experience, adaptation occurs, as indicated by a compensatory shift opposite to the prism deviation when subjects are asked to point straight-ahead after the prisms are removed (Redding et al. 2005). This remarkable perceptual adaptation seems to occur in the cerebellum (Luauté et al. 2009) and in regions of superior temporal cortex implicated in visuomotor action, rather than in occipital areas. This suggests adaptation is the result of recalibrating the mapping between visuomotor coordinate systems rather than recalibrating the visual input per se. It is worth noting that the transformations induced by prisms are distinct from the distortions found in macular degeneration patients, or those described in Figures 1 and 2. Prisms displace or invert the visual input with no loss of information and only very minor topographical distortion. Affine transformations such as these may be easier to decode from the retinotopically organized output of cortex.

Perhaps the most extreme example of higher visual areas compensating for V1 distortion might be circumstances where V1 is absent. In monkeys, visual training after V1 lesions restores the ability to detect and localize visual stimuli in the blind fields (Weiskrantz and Cowey 1963, Mohler and Wurtz 1977). However, in humans, attempts to restore conscious vision after V1 damage with visual restoration therapy have had limited success: reports indicate that the ‘restored’ conscious vision is far from normal, and restoration therapy has been shown to be ineffective in cortical blindness patients (for reviews see Pambakian and Kennard 1997, Melnick et al. 2016).

In summary, although higher level areas do seem to be able compensate for certain adult-onset distortions in V1, these abilities are constrained. One possibility is that there may be less flexibility in decoding distortions of the representation of retinotopic organization than there is for features that are more heavily influenced by developmental visual experience, such as disparity, direction, orientation tuning, and visuomotor calibration.

### 7.3. Why does plasticity vary across the visual hierarchy?

There are many possible reasons why the extent of cortical plasticity might vary across the hierarchy. One possibility is that this simply reflects the cumulative effects of plasticity at multiple preceding processing stages. Learning at each stage of processing presumably involves a selective reweighting of the connections from neurons that feed into that stage, with neurons best tuned for optimal performance given more weight (Saarinen and Levi 1995, Dosher and Lu 1998, Fine and Jacobs 2000). Given that the reweighting of neurons at each stage in the hierarchy presumably propagates to the next stage (and likely also influences lower stages due to cortical feedback) an increase in plasticity as a function of the number of preceding stages is not surprising. As mentioned above, significantly more subcortical processing occurs within somatosensory and auditory pathways than within visual pathways. Thus, A1 and S1 can be considered as ‘higher’ in their respective processing pathways than V1 is within the visual processing hierarchy. This may explain the informal observation that tactile and auditory tasks seem to show stronger perceptual learning effects than their vision ‘equivalents’.

A second, non-exclusive, possibility is that synaptic plasticity may genuinely vary across the visual hierarchy. There are a number of reasons why representations within early stages of visual processing may, by necessity, be resistant to plasticity in adulthood. For example, the representational demands of early visual cortex are unlikely to change substantially across the lifespan. Early visual areas represent the entire feature space (e.g. all orientations and spatial locations), although the sampling of that feature space is heavily dependent on childhood experience. Because this representation is complete and the low-level statistics of the environment do not change dramatically over the lifespan (i.e. we don’t suddenly experience new orientations in the same way as we experience new faces), there is little need for adult plasticity.

Indeed, allowing adult plasticity at early stages of processing might be expected to have deleterious consequences within later stages in the visual processing hierarchy. Any change in an early representation requires compensatory changes within all downstream mechanisms. Given that many complex objects are only experienced relatively infrequently, this compensation would presumably have to occur preemptively to experiencing those objects. Currently, there is no known neurobiological mechanism that could explain how this compensation could occur.

Finally, differences in plasticity across the cortical hierarchy may be the consequence of qualitative differences in how information is represented. As described above, early stage representations seem to reflect specific ‘features’ whereas representations in higher-level areas may occur at a distributed network level (Levy et al. 2004) that may be more amenable to experience-dependent change.

## 8. EFFECTS OF TRAINING IN VISUAL PROSTHESIS PATIENTS

Visual prosthesis and optogenetic technologies offer a unique platform to study cortical plasticity in adulthood. We have seen that local connections in adult cortex might be at least partially modifiable, but is this plasticity sufficient to make visual prostheses useful?

Previous experience with cochlear implants would suggest optimism. Adult-implanted cochlear implant users initially report extremely unnatural and incomprehensible perceptual experiences. Over remarkably short time periods, not only do subjects become better at important tasks such as speech recognition, but their perceptual experiences become more qualitatively ‘natural’. As described by one patient: *“It sounded like popcorn… every time someone said something it popped… as soon as I opened my mouth it popped… Now it sounds a little like Alice in Wonderland when she stands there in the tunnel talking. It is like more of a clanging sound now and it is very clear but still a bit lighter… more treble and like talking in a can.”* (Hallberg and Ringdahl 2004).

The abilities of visual prosthetic patients also improve with training. However there is little evidence that this is due to distortions becoming less perceptually apparent. Instead most improvements seem to be relatively task specific, and are the result of patients becoming better at interpreting distorted input.

One study did find reductions in visual field loss as measured with Goldmann perimetry after prosthetic implantation (Rizzo et al. 2015) for both implanted and non-implanted eyes. This may have been due to ‘reactivation’ of the visual system (though other phenomena such as Charles Bonnet syndrome (Cogan 1973) and the photophobia often observed after cataract surgery suggest that long-term deprivation leads to *upregulation* of deprived visual cortex). Alternatively, these results may reflect changes in behavioral response criteria (Shapiro and Johnson 1990). Another study found decreases in detection thresholds as a function of time since surgery (Castaldi et al. 2016). However subject performance on other relatively simple perceptual tasks, such as contrast sensitivity (Castaldi et al. 2016) and motion discrimination (Dorn et al. 2013, Castaldi et al. 2016) barely improved, even after extensive training over several months.

The largest perceptual improvements found in prosthetic users have generally been reported in tasks such as moving independently in space, locating a large bright square on a screen, and identifying large-print letters (∼40°) at above-chance levels (Chader et al. 2009, Zrenner et al. 2011, Humayun et al. 2012, da Cruz et al. 2013, Dorn et al. 2013, Stingl et al. 2013, Rizzo et al. 2015). After practice, single-letter recognition seems to take somewhere between a few seconds to three and a half minutes, enabling the best performing patients to read short words (da Cruz et al. 2013).

One possibility is that the perceptual ‘experience’ of prosthetic users will gradually improve over time, as seems to occur for cochlear implants, due to either a reduction in perceptual distortions or an increase in the perceptual discriminability of stimuli. Alternatively, patients may learn how to use distorted information to perform specific tasks. Dissociating these two kinds of learning, and understanding how they contribute to prosthetic vision, will require careful thought in future longitudinal studies of sight restoration patients.

Importantly, many assessments of the effects of training on visual prosthetic performance have focused on “closed” tasks under somewhat unrealistic contexts. This may lead to overly optimistic assessments of performance and the effects of training. An example of a strongly closed task is “is that an ‘A’ or a ‘B‘“. An example of a less closed task is “what letter is this?” An open task is “What is on the screen?” One difficulty with closed tasks, particularly those that mimic open tasks, is that they can give a highly misleading impression of performance. As in Figure 4, when subjects are told that the four items on the table include a plate, a cup, a napkin and a fork it is easy to see how subjects can successfully ‘find the fork’. Real dinner tables tend to be more cluttered and disorganized. An individual who can successfully find a fork among a preselected set of four items may not be able to find a candlestick on a cluttered dinner table. A second issue with closed tasks is that they are likely to result in perceptual learning that is highly specific to the particular context or task used (see Section 5.2 above). The literature on perceptual learning, described above, indicates very strongly that learning will almost always occur at the most specific level possible given the training task. Thus, care should always be taken when designing training protocols to ensure that improvements are likely to generalize to performance outside the laboratory or clinic.

**Figure 4.**
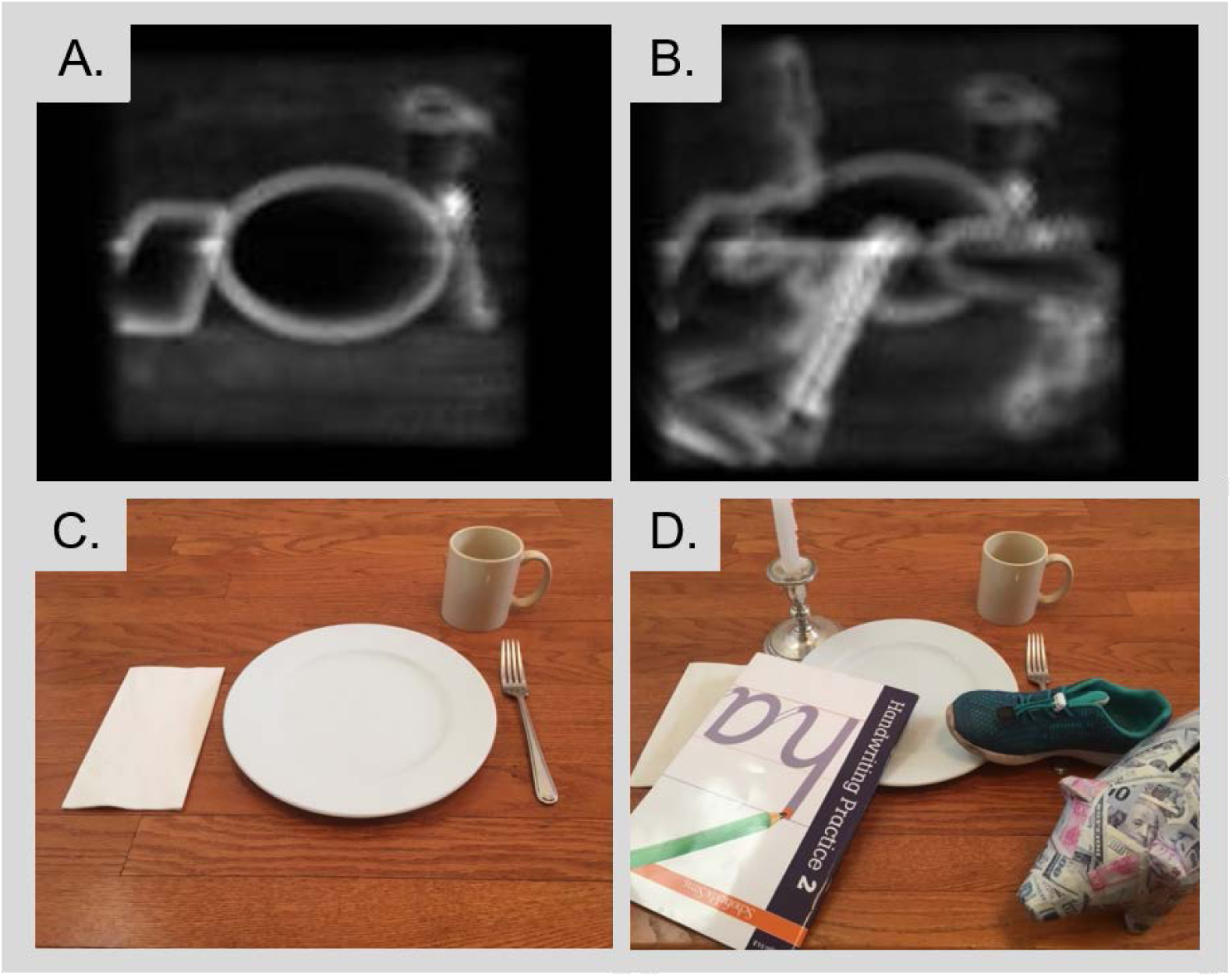
Two examples of a dining table under conditions of simulated prosthetic vision. Under ‘laboratory’ conditions the plate, cup, napkin and fork are easily differentiable. Real-world conditions for Dr. Fine’s dining table include multiple additional and unexpected objects. Finding the fork is challenging, even for an individual with normal vision.

## 9. REAWAKENING THE CRITICAL PERIOD

One exciting avenue for improving prosthetic implant outcomes may come from harnessing new methods for ‘reawakening the critical period’ (Bavelier et al. 2010, Werker and Hensch 2015) via pharmacological intervention.

Recent advances in neurobiology have provided an increasingly detailed picture of the cascade of events that control critical period plasticity. During early development, the synaptic neurochemical balance is shifted towards excitation, and synaptic connectivity is fluid. Synaptic rewiring includes physical pruning and homeostatic regrowth of synapses (controlled by the mediators tPA, TNFα, protein synthesis). Over time, responses to sensory input cause the promotion of inhibitory parvalbumin cell maturation and their increased GABA function through molecular triggers, such as orthodenticle homeobox 2 (Otx2), and brain-derived neurotrophic factor (BDNF, Huang et al. 1999). This shifts the excitation/inhibition balance towards a mature state of greater inhibition. The closure of the critical period is actively enforced by structural consolidation, including formation of a perineuronal extracellular matrix, and epigenetic brakes on plasticity (e.g., histone deacetylation) that silence the gene programs necessary for synaptic rewiring (Hensch 2005b, Takesian and Hensch 2013, Werker and Hensch 2015). This has led to a significant amount of research with the goal of ‘reawakening’ cortical plasticity via biochemical intervention that targets various aspects of this cascade.

In both animals and humans there is evidence that serotonin uptake inhibitors shift the excitatory inhibitory balance towards a state of greater excitation, and may enhance plasticity. In adult animal models fluoxetine results in reduced GABAergic inhibition and increased BDNF expression, which results in reopening of the critical period (Maya-Vetencourt et al. 2008, Bachatene et al. 2013), but a study in humans did not find that fluoxetine enhanced perceptual learning (Lagas et al. 2016). In humans the FDA-approved cholinesterase inhibitor donepezil seems to enhance perceptual learning, perhaps by shifting the excitation/inhibition balance towards the ‘immature’ state of greater excitation (Rokem et al. 2010, Rokem and Silver 2013, Chamoun et al. 2017). While shifting the neurochemical balance towards greater excitation may be necessary to ‘reopen’ the critical period, there is growing evidence that it is not sufficient (Hensch 2005a, Wong 2012, Takesian and Hensch 2013). Epigenetic brakes and structural limitations must also be overcome.

As far as epigenetic limitations are concerned, in humans, valproate has been successfully used to enhance auditory cortex plasticity for training in pitch perception in adult humans (Gervain et al. 2013), although these findings have yet to be replicated and might not generalize to the visual system. The use of valproate to target *cortical* plasticity is complicated by its adverse side effects (Sandberg et al. 2011, Sisk 2012, Bhalla et al. 2013) and a literature leaving it unclear whether its effect on *retinal* function in patients with retinal dystrophies is positive (Clemson et al. 2011, Iraha et al. 2016) or negative (Sisk 2012, Bhalla et al. 2013).

As far as structural limitations are concerned, homeobox protein Otx2 may eventually serve a critical role in reawakening the critical period. Disruption of Otx2 reduces parvalbumin (thereby influencing excitatory/inhibitory balance) and reduces perineuronal net expression (thereby reducing structural limitations on plasticity). In animal models disruption of Ox2 has proved sufficient to reopen plasticity in adult mice (Beurdeley et al. 2012, Spatazza et al. 2013). However, pharmacological enhancement of OTX2 raises serious safety concerns given that over-expression of OXT2 is associated with a variety of cancers (Di et al. 2005, Nagel et al. 2015).

In conclusion, as we develop a fuller understanding of the triggers, modulators and brakes that control the sensitive period, it appears increasingly likely that ‘reawakening plasticity’ in cortex is likely to require simultaneously targeting multiple parts of the cascade: it is not adequate to reduce structural brakes on synaptic rewiring if the excitatory-inhibitory balance is unsuitable for new connection patterns to establish, and vice versa. However, as described above, there are numerous FDA-approved drugs that seem likely to modestly enhance plasticity. Any reader over the age of forty is well aware that adult plasticity continues to decline with age. Given that most prosthetic implant recipients are likely to be elderly, even modest improvements in the capacity for rehabilitative plasticity might greatly enhance their ability to make functional use of their devices.

## 10. CONCLUSIONS

Visual prostheses and optogenetic methods for restoring retinal function offer a unique platform for studying cortical plasticity in adults that is likely to increase in translational importance as the field progresses. The literature described in this review strongly suggests that there exists a “tolerance envelope” of adequate sensory information which any successful prosthetic device must match. If distortions or the amount of information loss fall within that envelope, it seems plausible that cortical plasticity can successfully compensate. However, it must be recognized that cortical plasticity is finite and cannot compensate for distortions or information loss that falls outside that envelope.

Thus, it is critical to understand which visual distortions can be compensated for via cortical plasticity, which cannot, and what role training or pharmacological intervention can play. Engineers need to focus their energies on developing technologies that fall within the sensory ‘tolerance envelope’. This will require a shift from thinking of the brain as having ‘pluripotent plasticity’ towards a more nuanced understanding of which aspects of visual perception are adaptable, and under which conditions. It is not clear whether spatial distortions can be remapped within early visual areas, but higher level areas may be surprisingly competent at decoding a distorted world. However, learning to decode a distorted world must be done in the context of generalizable learning; this will require training across a wide range of stimulus sets or navigational environments, while maintaining high performance levels. Understanding these principles of effective and generalizable training will be critical for developing rehabilitation strategies that can efficiently help patients make best use of their implants. In the context of prosthetic vision, this may require increased use of virtual reality environments.

Finally low-level visual plasticity seems to be limited compared to other cognitive and sensory domains. This provides a significant technical challenge for engineers, and any way of enhancing low-level plasticity is likely to be extremely beneficial for rehabilitative training. In the short-term, gamification may provide significant benefits. In the longer term pharmacological intervention provides an exciting potential opportunity.

What is most exciting about this relationship between neural prosthesis development and basic neuroscience is that the flow of information will be reciprocal. The next decade is likely to yield fundamental insights within an important, but relatively poorly understood field of research – human adult plasticity. Perhaps the capacity that most differentiates humans from other animals is our capacity to continuously learn new perceptual and cognitive skills throughout our lives. Visual prosthesis patients are likely to provide a unique opportunity to gain insights into the fundamental principles that underlie these abilities.

## 11. ACKNOWLEDGMENTS

Supported by the Washington Research Foundation Funds for Innovation in Neuroengineering and Data-Intensive Discovery (M.B.), by a grant from the Gordon & Betty Moore Foundation and the Alfred P. Sloan Foundation to the University of Washington eScience Institute Data Science Environment (M.B. and A.R.), and by the National Institutes of Health (NEI EY-12925 to G.M.B., EY-014645 to I.F.).

